# S2 from Equine infectious anemia virus is an infectivity factor which counteracts the retroviral inhibitors SERINC5 and SERINC3

**DOI:** 10.1101/065078

**Authors:** Ajit Chande, Cristiana Cuccurullo, Annachiara Rosa, Serena Ziglio, Susan Carpenter, Massimo Pizzato

## Abstract

The lentivirus equine infectious anemia virus (EIAV) encodes S2, a pathogenic determinant important for virus replication and disease progression in horses. No molecular function has yet been linked to this accessory protein. We now report that S2 can replace the activity of Nef on HIV-1 infectivity, being required to antagonize the inhibitory activity of SERINC proteins on Nef-defective HIV-1. Similar to Nef, S2 excludes SERINC5 from virus particles and requires an ExxxLL motif predicted to recruit the clathrin adaptor AP2. Accordingly, a functional endocytic machinery is essential for S2-mediated infectivity enhancement, which is impaired by inhibitors of clathrin-mediated endocytosis. In addition to retargeting SERINC5 to a late endosomal compartment, S2 promotes the host factor degradation. Emphasizing the similarity with Nef, we show that S2 is myristoylated and, compatible with a crucial role of the post-translational modification, its N-terminal glycine is required for the anti-SERINC5 activity.

EIAV-derived vectors devoid of S2 are less susceptible than HIV-1 to the inhibitory effect of both human and equine SERINC5. We then identified the envelope glycoprotein of EIAV as a determinant which also modulates retrovirus susceptibility to SERINC5, indicating a bi-modular ability of the equine lentivirus to counteract the host factor.

S2 shares no sequence homology with other retroviral factors known to counteract SERINC5. Adding to primate lentivirus Nef and gammaretrovirus glycoGag, the accessory protein from EIAV makes another example of a retroviral virulence determinant which independently evolved SERINC5-antagonizing activity. SERINC5 therefore plays a critical role for the interaction of the host with diverse retrovirus pathogens.

**Significance Statement:** SERINC5 and SERINC3 are recently discovered cellular inhibitors of retroviruses, which are incorporated into virus particles and impair their ability to propagate the infection to target cells. Only two groups of viruses (represented by HIV-1 and MLV) have so far been identified to have evolved the ability of counteracting SERINC inhibition. We now discovered that Equine infectious anemia virus, which causes a debilitating disease in horses, also acquired the ability to protect the virus particle from inhibition by SERINC5 and SERINC3, using its small protein S2. The evidence that three different retroviruses have independently evolved the ability to elude inhibition bySERINC5 and SERINC3 indicates that these cellular factors play a fundamental role against various retrovirus pathogens.

## Introduction

During the process of adaptation to a host, viruses acquire the ability to exploit the cell and, at the same time, to overcome barriers that interfere with their replication. Several cellular factors, also named “restriction factors”, have been identified during the past 30 years, capable of inhibiting different steps in the retrovirus life cycle and signaling an ongoing pathogen invasion(1). Retroviruses have however evolved the ability to overcome such cellular inhibitors either by escaping their recognition or by actively antagonizing them. Some retroviruses can count on an increased plasticity conferred by the complexity of their genome, exploiting alternative splicing to encode accessory proteins, which have often evolved the ability to antagonize intrinsic host defenses (2). Nef from primate lentiviruses and glycoGag from gammaretroviruses were found to exert a convergent activity(3), by similarly promoting the infectivity of HIV and MLV. The molecular basis for such functional similarity was revealed recently, as they were both found capable of counteracting SERINC5, a cellular multipass transmembrane protein highly expressed in lymphoid and myeloid tissues(4, 5). SERINC5 inhibits an early stage of retrovirus infection of target cells, following its incorporation into virions. Both Nef and glycoGag antagonize SERINC5 by promoting its intracellular relocalization *via* a clathrin-dependent mechanism, and preventing its incorporation into virions. Human SERINC5 inhibits efficiently the divergent retroviruses HIV-1 and MLV, indicating low species specific barriers. Given that Nef and glycoGag have evolved the ability to counteract SERINC5 independently in primate lentiviruses and gammaretroviruses, we sought to investigate whether the selective pressure imposed by this host factor has affected the evolution of other retroviruses. SERINC5 being highly expressed in blood-derived cells, we investigated whether another blood-tropic retrovirus, Equine infectious anemia virus (EIAV), could have also evolved a Nef-like infectivity factor. EIAV is a myeloid-tropic lentivirus which causes anemia and thrombocytopenia in horses worldwide, and chronicizes into an “inapparent” asymptomatic carrier stage, critical for virus dissemination to uninfected horses(6). EIAV is a complex retrovirus, which expresses the 7KDa auxiliary protein S2 from a doubly spliced mRNA(7). S2 has no homology to other known proteins and remains an orphan with unknown molecular activity. Intriguingly, mutant EIAV lacking a functional S2 ORF resembles *in vivo* phenotypically a Nef-defective HIV-1. In fact, while there is no evidence for a requirement of S2 for EIAV replication *in vitro*, its requirement as pathogenic factor *in vivo* has been well established, since infection of animals with a virus lacking a functional S2 is associated with low viral load and the absence of clinical symptoms(8–10).

Here we report that S2 has evolved the ability to antagonize SERINC5 and SERINC3, providing further indication, along with Nef and glycoGag, that the recently discovered host restriction factor plays a crucial role in the biology of retroviruses and has shaped their evolution.

### S2 rescues the infectivity of Nef-defective HIV-1

We first investigated whether S2 could functionally replace the activity of Nef on HIV-1 infectivity. S2 ORF cloned from the pSPEIAV19 strain(11) was inserted into an *env*-defective HIV-1_NL4-3_ molecular clone in place of *nef* (Fig. 1A). The resulting construct was used to produce HIV-1 limited to a single replication cycle by transfecting JTAg cells. HIV-1 encoding S2 was 6 fold more infectious than *nef*-defective HIV-1_NL4-3_, indicating that S2 can restore most of the infectivity lost in the absence of Nef. In order to monitor the expression of S2 by Western blotting and immunofluorescence, the S2 coding squence was optimized based on human codon usage and fused to an HA tag. While the N-terminally tagged protein was poorly detected by Western blotting (and inactive on HIV-1 infectivity), the HA tag at the C-terminus was detected and its activity was similar to the untagged protein expressed from the native ORF (Fig. S1). S2-HA was therefore used in subsequents experiments. When S2-HA was expressed in *trans*, it restored the infectivity of the Nef-defective HIV-1 to a level similar to the WT counterpart (Fig. 1*B*) while, at the same time, it did not significantly alter HIV-1 infectivity in the presence of Nef, indicating that the activity of S2 and Nef are complementary rather than additive. The ability of S2 to restore the infectivity of Nef-defective HIV-1 was further confirmed using a different S2 allele, derived from the highly pathogenic EIAV strain Wyoming (Fig. S2).

**Figure 1.**
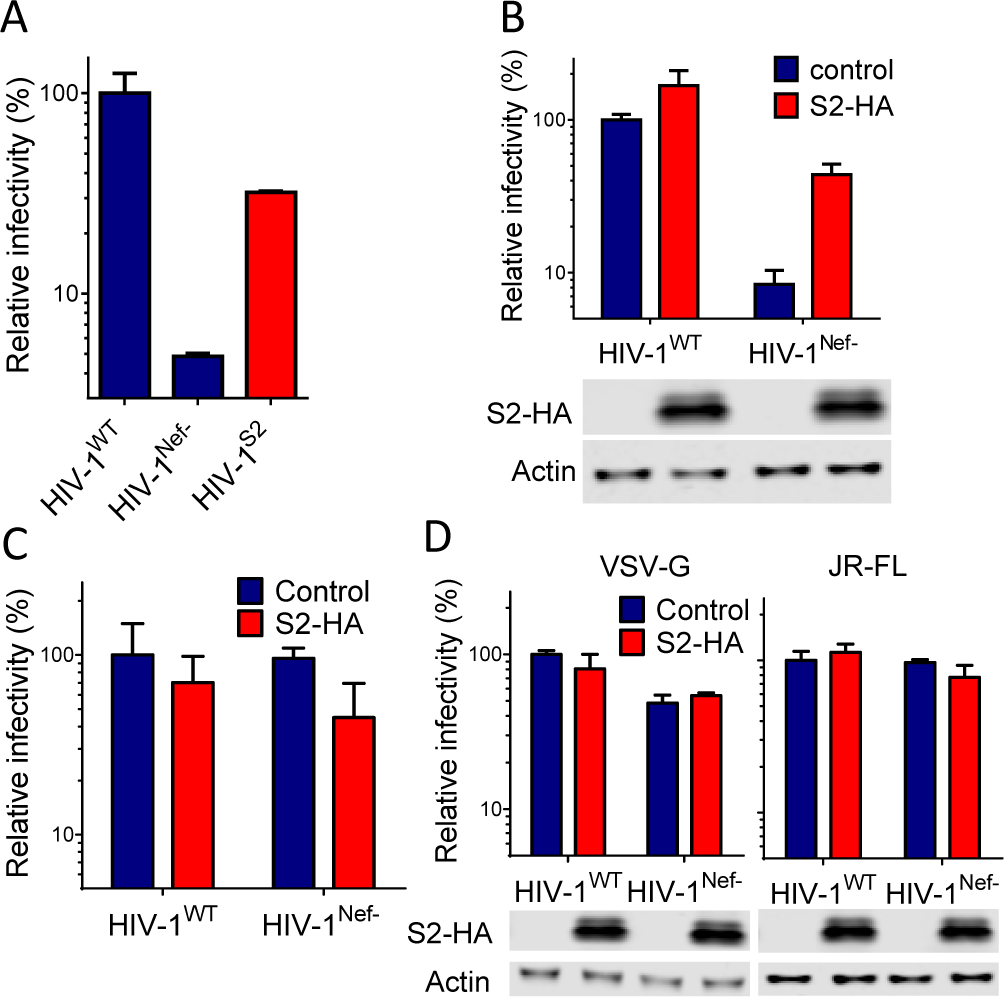
EIAV S2 and HIV-1 Nef are functionally similar infectivity-promoting factors. S2 repairs the defective infectivity of Nef-defective single cycle HIV-1_NL4-3_ produced in JTAg cells (A and B). S2 was expressed in place of Nef within the HIV-1_NL4-3_ provirus (A) or *in trans*, fused to an HA tag at the C-terminus (B). C and D: the effect of S2 on infectivity is subjected to the same requirements of the effect of Nef, since S2 does not affect the infectivity of HIV-1_NL4-3_ derived from CEMx174 (E) or HIV-1_NL4-3_ pseudotyped with VSV-G or JRFL Env (D). Relative infectivity is expressed as % of the infectivity of wt HIV-1_NL4-3_ in the absence of S2 expression. Error bars represent standard deviation of the mean calculated from quadruplicate determinations. B and D include Western blots detection of S2-HA and beta-actin in lysates of virus producing cells.

The effect of Nef on HIV-1 infectivity has distinctive features, since it depends on the Env glycoprotein and the producer cell-type (4). We therefore investigated whether the effect of S2 shares the same features. As established previously, HIV-1 produced from CEMX174 cells does not require Nef for producing optimally infectious virus. Similarly, S2 had no effect on the infectivity of HIV-1 derived from CEMX174 cells (Fig. 1C). HIV-1 pseudotyped with VSV-G or with the Env glycoprotein derived from HIV-1_JRFL_ is not responsive to Nef (Fig. 1D). Similarly, S2 expression in JTAg producer cells did not alter the infectivity of Nef-defective HIV-1 progeny virus pseudotyped with VSV-G or HIV-1_JR-FL_ Env (Fig. 1D). Nef and S2 therefore are subjected to similar cell-type and Env dependence, further highlighting a functional similarity.

### S2 counteracts SERINC5 and SERINC3

We have recently discovered that the host factors SERINC5 and SERINC3 are inhibitors of HIV-1 and MLV and that they are counteracted by Nef and glycoGag respectively(4). We therefore investigated whether S2 acts by antagonizing the same restriction factors antagonized by Nef.

To this end we first investigated whether the ability of S2 to increase HIV-1 infectivity in JTAg cells requires both SERINC5 and SERINC3. CRISPR/Cas9 was used to generate JTAg cells lacking either SERINC5 alone (JTag^SERINC5 KO^) or SERINC5 in combination with SERINC3 (JTag^SERINC5/3 KO^). Nef-defective HIV-1 derived from wt JTAg cells is characterized by 10-fold lower infectivity than virus derived from JTag^SERINC5/3 KO^ and 3-fold lower infectivity than virus derived from JTag^SERINC5 KO^ (Fig. 2A). This is consistent with the higher inhibitory activity of SERINC5 than that of SERINC3. Compatible with the ability to counteract both restriction factors, expression of S2 resulted in a 10-fold increase of the infectivity of HIV-1 produced from wt cells and only a 3-fold increase of the infectivity of virus derived from JTag^SERINC5 KO^. In contrast, S2 did not alter the infectivity of HIV-1 derived from JTag^SERINC5/3 KO^(Fig. 2A). The effects of S2 on HIV-1 was paralleled by the effect of Nef, which increased 12-fold, indicating that both retroviral accessory proteins enhance HIV-1 infectivity by similarly counteracting SERINC5 and SERINC3.

**Figure 2.**
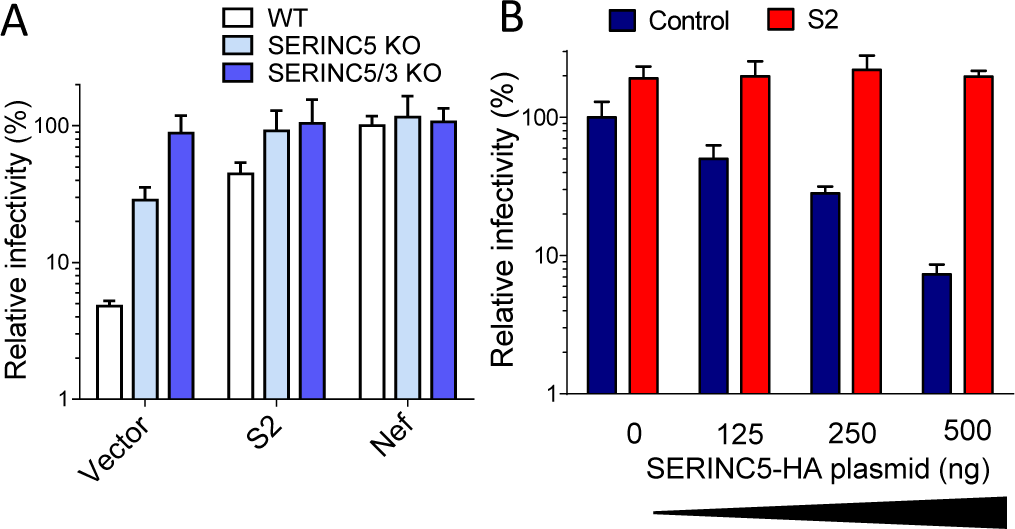
S2 counteracts inhibition of HIV-1 by SERINC5 and SERINC3. A, the effect of S2 on the infectivity of Nef-defective HIV-1_NL4-3_ produced in JTAg cells requires the endogenously expressed SERINC5 and SERINC3. Nef-defective HIV-1^NL4-3^ produced in JTAg cells knocked out for SERINC5 or SERINC5 in combination with SERINC3. B, S2 preserves the infectivity of Nef-defective HIV-1_NL4-3_ from inhibition by increasing amount of PBJ6-SERINC5-HA plasmid transfected in producer HEK293T cells. Relative infectivity is expressed as % of the infectivity of Nef-defective HIV-1_NL4-3_ in the absence of S2 expression. Error bars represent standard deviation of the mean calculated from quadruplicate determinations.

To further confirm the ability of S2 to antagonize SERINC5, the host factor was ectopically expressed in HEK293T cells during virus production. Increasing amounts of SERINC5 expressing plasmid inhibited Nef-defective HIV-1 infectivity from 2 to 50-fold in the absence of S2-HA (Fig. 2B). In contrast, virus infectivity remained unchanged when S2 was co-expressed in producer cells, irrespectively of the amount of SERINC5-HA plasmid transfected.

The EIAV accessory protein therefore promotes virus infectivity by counteracting SERINC5 and SERINC3.

### S2 prevents SERINC5 incorporation into virus particles

We have previously established that SERINC5 is efficiently incorporated into retroviral particles and that HIV-1 Nef and MLV glycoGag act in producer cells by promoting SERINC5 exclusion from virions. We investigated whether S2 acts in a similar way. Nef-defective HIV-1 particles were produced in HEK293T cells transfected to express HIV-1, SERINC5-HA and S2-HA. As revealed by Western blotting, while SERINC5-HA was readily detected in Nef-defective virions, expression of S2-HA drastically reduced its incorporation as the protein was detected at a level similar to background (Fig. 3A). S2, like Nef and glycoGag, therefore acts by excluding the host factor from virus particles.

**Figure 3.**
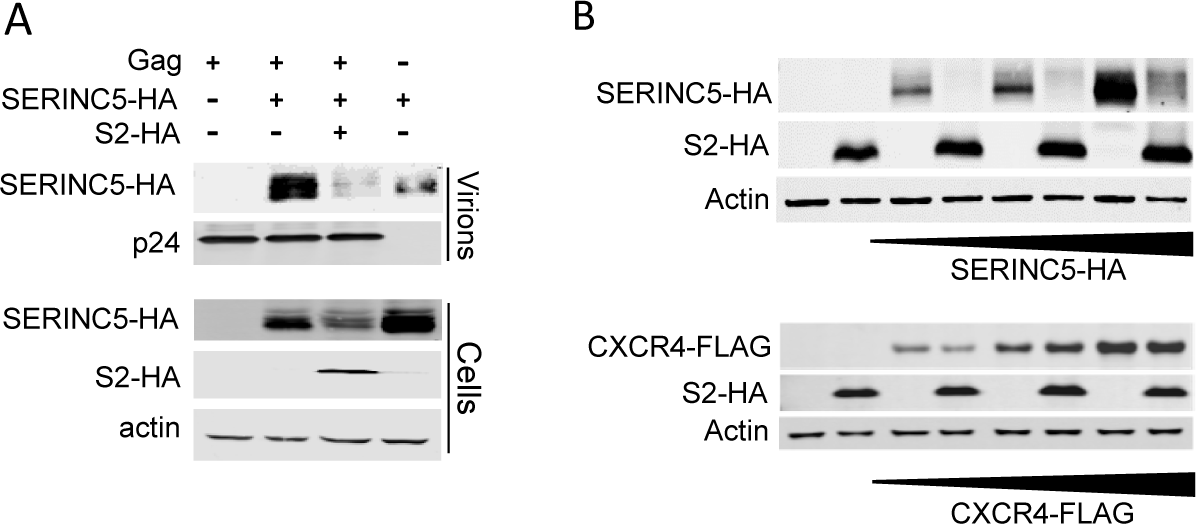
S2 prevents virion incorporation of SERINC5. S2 expression in HEK293T cells producing Nef-defective HIV-1_NL4-3_ prevents virion incorporation of SERINC5-HA. Western blotting of virion pellets and lysates of respective producer cells. B, S2-HA alters steady state expression level of SERINC5-HA but not CXCR4-FLAG. Western blotting of lysates from HEK293T cells that were co-transfected with a plasmid encoding S2-HA and increasing amounts of SERINC5-HA or CXCR4-FLAG encoding vectors.

We observed that the steady-state level of SERINC5 in the lysate of virus producing cells was reduced when S2 was expressed (Fig. 3A), indicating a possible effect of S2 on SERINC5 stability. To further investigate this possibility, the effect of S2 was tested with an increasing amount of SERINC5-HA expressing plasmid and a constant amount of vector expressing S2-HA in HEK293T cells. S2 co-expression resulted in a marked decrease of SERINC5-HA in cell lysates, irrespectively of the amount of plasmid transfected (Fig. 3B). To investigate whether this effect was specific for SERINC5, the effect of S2 on the steady state level of CXCR4, another multipass transmembrane protein, was also tested. Contrary to the effect observed on SERINC5-HA, the steady state level of FLAG-tagged CXCR4 was not affected by S2 (Fig. 3B), irrespectively of the amount of the chemokine receptor being expressed. This indicates that the effect of S2 on the steady state level of SERINC5 is selective.

### S2 requires clathrin-dependent vesicular trafficking and a conserved putative di-leucine sorting motif to counteract SERINC5

Both Nef and glycoGag cause a dramatic relocalization of SERINC5 from the plasma membrane into a perinuclear Rab7-positive late endosomal compartment(4). We investigated whether S2 acts in a similar way on SERINC5 intracellular distribution. When SERINC5-GFP was expressed by transfecting JTAg cells, together with a control protein (RFP), its localization was almost exclusively restricted to the plasma membrane (Fig. 4A). In contrast, when a plasmid encoding S2-HA was co-transfected in place of RFP, the host factor readily relocalized into a perinuclear compartment and only partly co-localized with S2-HA. Co-transfection with a plasmid expressing Rab7-RFP revealed that the accumulation of SERINC5-GFP by S2 occurs in a compartment that is labelled with red fluorescence (Fig. 4B). This indicates that, similarly to Nef and glycoGag, S2 expression results in SERINC5 relocalization from the cell surface to the late endosomal compartment, suggesting the involvement of endocytosis.

**Figure 4.**
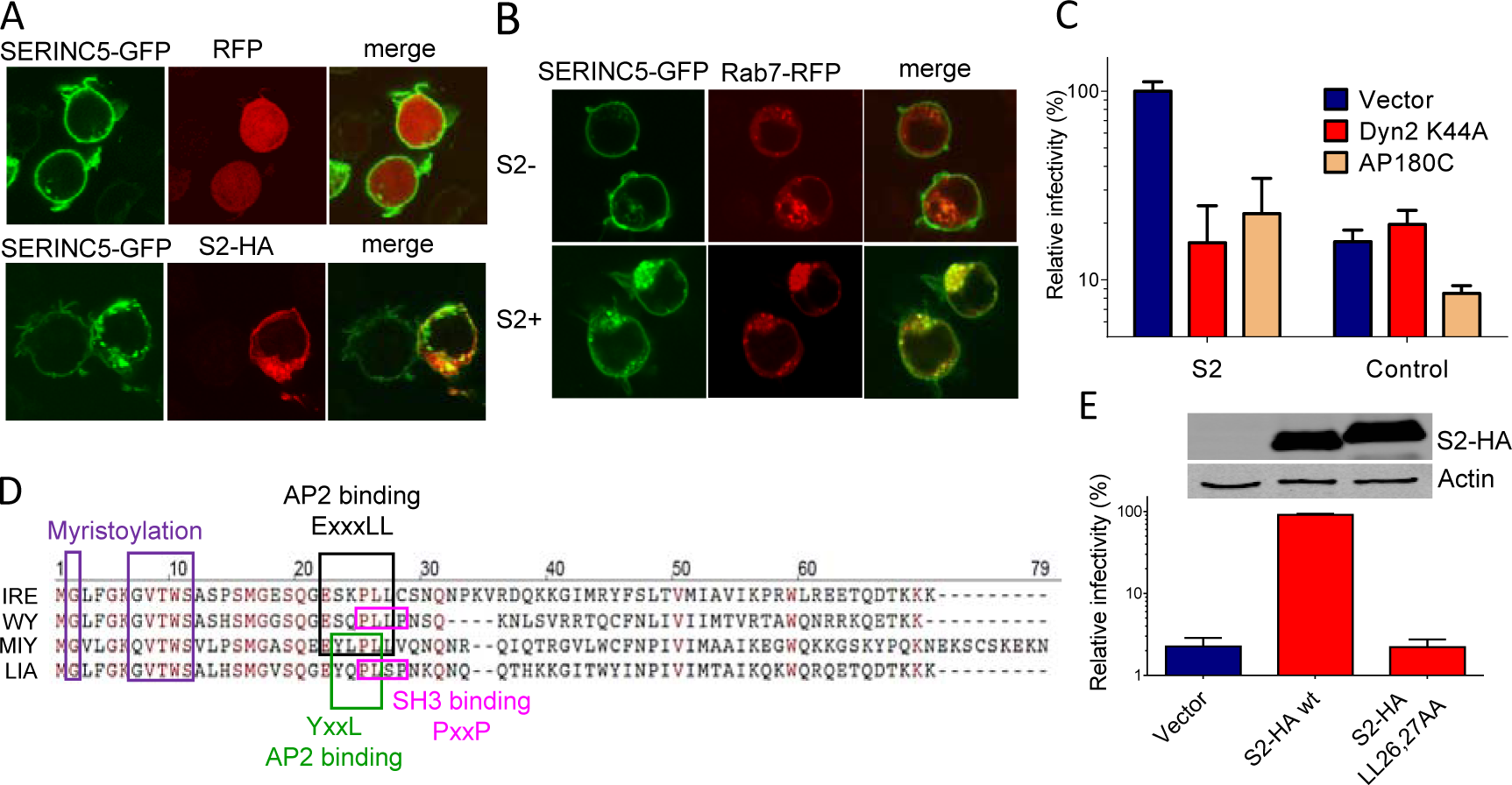
Counteraction of SERINC5 by S2 requires clathrin-dependent endocytosis and a conserved putative AP2 binding motif. A and B, S2 causes a relocalization of SERINC5-GFP from the plasma membrane to a Rab7-labelled compartment. Fluorescence confocal imaging of JTAg cells transfected to express the indicated proteins. C, the ability of S2, expressed in *trans*, to counteract SERINC5 is impaired by the clathrin-dependent endocytosis inhibitors Dyn2K44A and AP180-C in HEK293T cells transfected to produce Nef-defective HIV-1_NL4-3_. D, ClustalW alignment of S2 amino acid sequences derived from 4 isolates representing the four major EIAV groups: EIAV_wy_ (WY, accession AAC03764) EIAV _IRE_(IRE, accession AFW99166), EIAV_LIA_ (LIA, Accessio AAK21109) and EIAV_MIY_ (MIY, Accession AFV61766). Putative functional motifs are boxed and color-coded. D, integrity of the ExxxLL sequence is required for S2 counteracting activity of SERINC5 on Nef-defective HIV-1_NL4-3_. Virus produced by transfecting HEK293T cells expressing wild-type or mutant S2-HA. Western blotting show expression of S2-HA and beta-actin in lysates of producer cells. Infectivity in C and D is expressed as percentage relative to Nef-defective HIV-1_NL4-3_ produced in the presence of wt S2-HA and in the absence of inhibitors. Error bars represent standard deviation of the mean calculated from quadruplicate determinations.

The role of vesicular trafficking was further explored by assessing the effects of inhibiting clathrin-mediated coated pits formation on the ability of S2 to counteract SERINC5. Vectors encoding transdominant-negative dynamin 2 (Dyn2K44A), which inhibits vesicle constriction, and the C-terminus fragment of the clathrin adaptor AP180 (AP180C), which more specifically interferes with clathrin-dependent invagination of the cell surface (12), specifically the ability of the EIAV accessory protein to promote progeny virus infectivity (Fig. 4C), indicating that S2 engages the intravesicular trafficking machinery and requires clathrin-mediated endocytosis to counteract SERINC5.

A putative similarity between S2 and Nef is based on the phenotype of S2-defective EIAV *in vivo* and on the presence of motifs within the S2 amino acid sequence which can also be observed in the primate lentiviral protein(7–10, 13). These includes a possible myristoylation signal and a SH3 binding motif (PxxP). In addition, in S2 from different isolates we have observed the presence of either an ExxxLL or alternatively an YxxL putative AP2-binding motifs (Fig. 4D) similar to those found in Nef and glycoGag (14)(15). To verify whether ExxxLL is required for the effect of S2 on infectivity, the two leucine residues were mutated into alanine and the ability of the resulting mutant protein to counteract SERINC5 was tested. While the double mutation did not alter the expression level of the protein (Fig. 4E), it abolished its ability to promote the infectivity of the virus in the presence of SERINC5, indicating that the ExxxLL motif plays an important role for the activity of S2. In contrast, a PxxP motif, which could act as an SH3-binding domain, is dispensable for the activity of S2 on SERINC5. (Fig. S3)

### S2 is myristoylated

Both Nef and glycoGag associate with the plasma membrane, due to Nef being N-terminally myristoylated and glycoGag being an integral type-II transmembrane protein. It has previously been suggested that S2, like Nef, could be myristoylated(7, 13, 16), based on the presence of a possible canonical myristoylation signal located 5 residues from the N-terminus including Gly^7^ and Ser^11^ (Fig. 4D). However, experimental evidence confirming this possibility has never been reported. Since myristoylation occurs always at the N-terminal glycine, we investigated the importance of Gly^2^ for the ability of S2 to counteract SERINC5. An S2 mutant protein in which the N-terminal glycine was replaced with alanine (S2-G2A) was generated for this purpose. Mutant S2 G2A-HA was expressed in HEK293T producing HIV-1 in the presence of SERINC5-HA. While wt S2-HA rescued Nef-defective HIV-1 infectivity 10-fold in the presence of SERINC5 (Fig. 5A), the mutant protein failed to counteract the host factor, despite being detected in cell lysates at a similar level as the WT protein (Fig. 5A). The N-terminal glycine plays therefore a crucial role for the ability of S2 to counteract SERINC5, consistent with the possible involvement of myristoylation.

**Figure 5.**
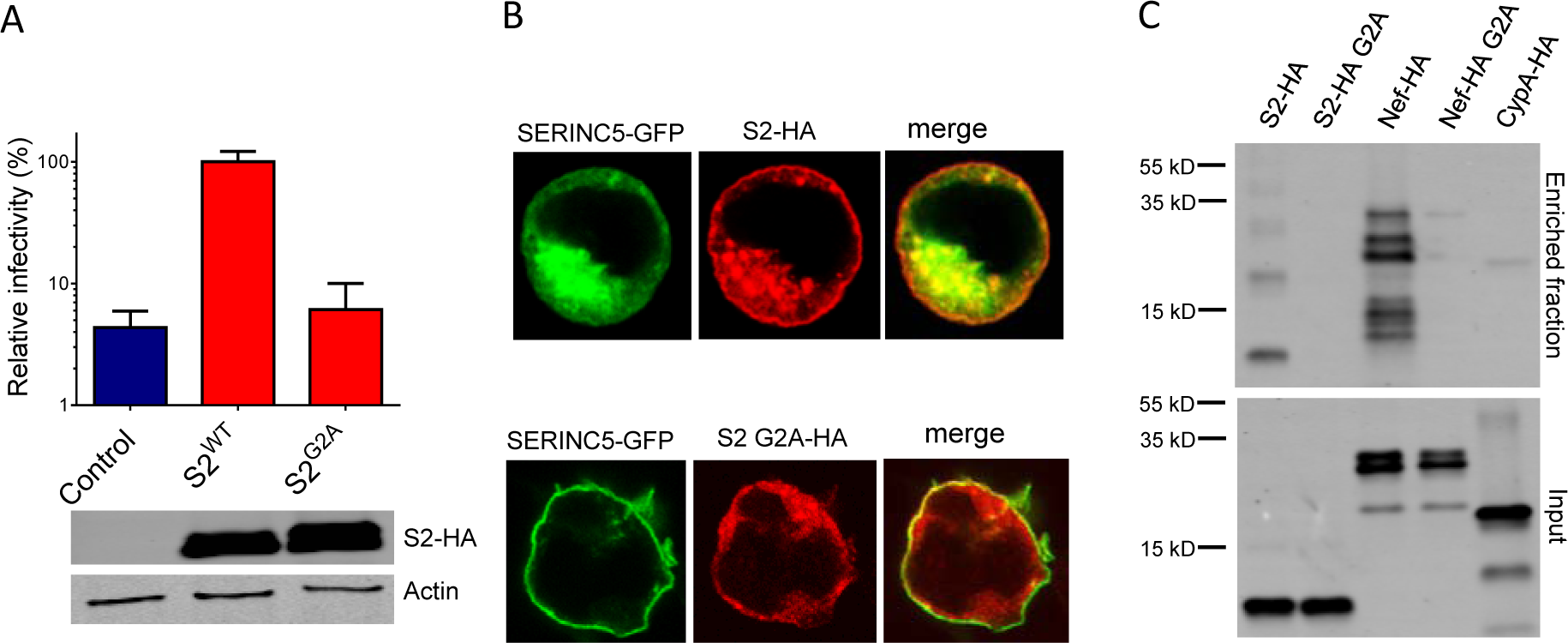
The role of N-terminal glycine for S2 activity. A, Gly^2^ is required for S2-HA counteracting activity of SERINC5 on Nef-defective HIV-1_NL4-3_. Virus was produced by transfecting HEK293T to express wild-type or mutant S2-HA. Western blotting shows expression of S2-HA and beta-actin in lysates of producer cells. Infectivity is expressed as percentage relative to the infectivity of Nef-defective HIV-1_NL4-3_ produced in the presence of wt S2-HA. Error bars represent standard deviation of the mean calculated from quadruplicate determinations. B, Gly^2^ is required for S2-HA localization at the cell membrane. immunofluorescence confocal imaging of JTAg cells transfected to express the indicated. C, S2 is myristoylated. Western blotting to detect HA-tagged proteins in the myrstic azide-enriched fraction (left) and in whole lysates (right) derived from HEK293T cells labelled with myrstic azide.

To directly assess *in vivo* whether S2 can be myristoylated, cells expressing S2-HA were labelled with azide-conjugated myristic acid and the presence of S2-HA in the azide-labelled enriched fraction of cell lysates was then probed by Western blotting using an anti-HA antibody. WT S2-HA (Fig. 5B) was readily detected in the enriched fraction, while only a signal indistinguishable from background was detected from samples expressing S2-HA lacking the N-terminal glycine (S2 G2A). Similarly, Nef-HA, but not NefG2A, was readily visible in the enriched fractions, while a non-myristoylated control protein (CypA-HA), produced no signal above background. These results indicate that EIAV S2 can be specifically targeted by the myristoyl transferase.

N-terminal myristoylation is instrumental for association with cellular membranes. Accordingly, as already observed in Fig. 4A, in transfected JTAg cells, wt S2-HA accumulates in vesicular structures partly localizing together with SERINC5-GFP, and at the cell periphery (Fig. 5C). However, the G2A S2 mutant, which failed to relocalize SERINC5, was instead distributed more diffusely throughout the cytoplasm. Altogether, these results suggest that the N-terminal glycine is crucial for S2 intracellular distribution and for its localization with vesicles and with the plasma membrane, and support a role of S2 myristoylation for the activity on infectivity.

### Sensitivity to SERINC5 is also modulated by EIAV Env

After having established the ability of S2 to counteract SERINC5 using HIV-1, the role of S2 was further investigated in the context of EIAV retrovirus particles.

Recombinant EIAV vector particles were produced by transfecting HEK293T in the presence and absence of ectopically expressed human SERINC5-HA. Infectivity of progeny recombinant particles was quantified by transducing TZM-bl cells engineered to stably express equine ELR1, the EIAV receptor(17). In the absence of S2, EIAV vector particles carrying an EIAV Env were inhibited by SERINC5 only 3 fold (Fig. 6A, left), indicating that EIAV retrovirus particles are less sensitive to the host factor than HIV-1 (Fig. 6A, right). However, in the presence of S2, SERINC5 did not affect EIAV vectors infectivity, confirming the ability of the accessory protein to counteract the inhibition exerted by the host factor.

**Figure 6.**
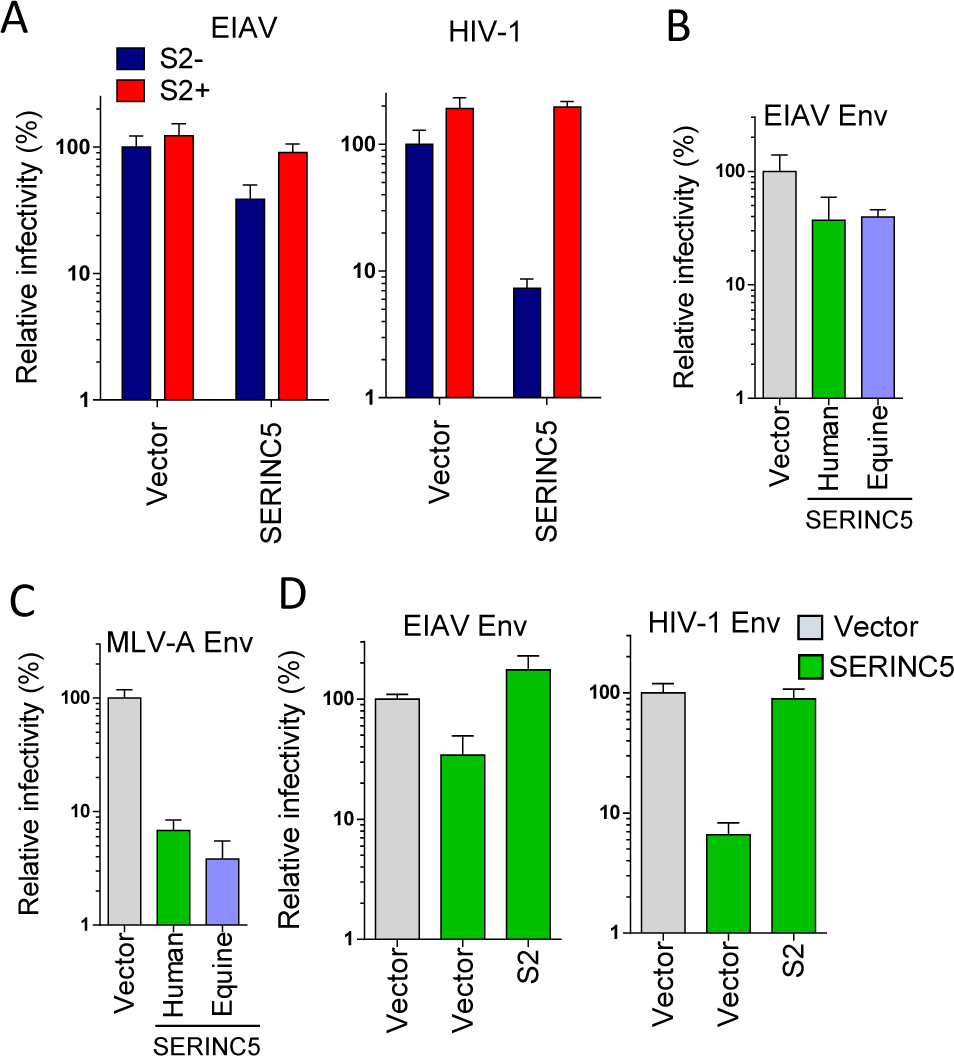
EIAV Env confers virus particles partial resistance to SERINC5. A, EIAV is less susceptible than HIV-1 to SERINC5 inhibition. Infectivity of S2-defective EIAV (left) and Nef-defective HIV-1 (right) virion particles by SERINC5 and counteraction by S2. B, S2-defective EIAV is equally sensitive to human and equine SERINC5. C, S2-defective EIAV pseudotyping with MLV-A Env confers increased susceptibility to both human and equine SERINC5. D, HIV-1 is less susceptible to SERINC5 when pseudotyped with EIAV Env than HIV-1 Env in the absence of S2. All panels show infectivity of single cycle EIAV and HIV-1 virus particles produced from HEK293T cells transfected to express also SERINC5 and S2 as indicated. Infectivity is expressed as percentage relative to the infectivity of Nef-defective HIV-1_NL4-3_ or S2-defective EIAV produced in the in the absence of SERINC5 overexpression. Error bars represent standard deviation of the mean calculated from quadruplicate determinations.

Given that inhibition of EIAV by human SERINC5 is weak compared with HIV-1, we investigated whether the magnitude is species-specific and therefore whether the equine retrovirus would be more sensitive to equine SERINC5. When human and equine SERINC5 were expressed in HEK293T producing EIAV vector particles, a similar magnitude of inhibition was observed irrespectively of the ortholog used (Fig. 6B), therefore ruling out a species-specific anti-retroviral activity. The EIAV vectors used are therefore intrinsically less sensitive to SERINC5, irrespectively of S2 expression.

The envelope glycoprotein is a crucial determinant of retrovirus particles susceptibility to SERINC5 (Fig. 1D). We therefore investigated whether the lower sensitivity of EIAV virions to SERINC5 is a feature associated with the viral core or depends on the Env glycoprotein. To discriminate between these two possibilities, we investigated the ability of SERINC5 to inhibit EIAV pseudotypes. After unsuccessfully attempting to generate infectious EIAV pseudotypes bearing HIV-1 Env, MLV-A Env was used instead to study EIAV vector particles carrying an Env glycoprotein known to render HIV-1 fully sensitive to SERINC5(4). Expression of either human or equine SERINC5 in producer cells resulted in a 10-20 fold defect of infectivity of the EIAV/MLV-A hybrid particles (Fig. 6C), indicating that when pseudotyped with a SERINC5-susceptible Env, EIAV virions also become fully susceptible to the host factor. This demonstrates that EIAV cores are not inherently resistant to SERINC5, but rather that Env plays a crucial role in modulating sensitivity to SERINC5. We therefore assessed the ability of EIAV Env to alter HIV-1 sensitivity to SERINC5, by studying inhibition of HIV-1 virion particles bearing EIAV Env. While Nef-defective HIV-1 carrying an HIV-1_HXB2_ Env was inhibited 15 fold by SERINC5, the same virion particles carrying EIAV Env were only affected 3-fold by the host factor (Fig. 6D), therefore mirroring the effect observed on EIAV vector particles (Fig. 6A). EIAV Env therefore confers partial SERINC5 resistance to retrovirus particles. Confirming its anti-SERINC5 activity, expression of S2 counteracted the effect of SERINC5, irrespectively of the Env glycoprotein used (Fig. 6C).

## Discussion

In this study, we have established that S2 from EIAV is an infectivity factor functionally related to Nef of primate lentiviruses and to glycosylated Gag of gammaretroviruses. S2 therefore provides a third example of a retroviral factor capable of promoting retrovirus infectivity by counteracting the SERINC family of proteins. Remarkably, the three retroviral factors share no sequence homology and originate from distinct and unrelated genetic regions of the retrovirus genome, indicating that the ability to counteract SERINC5 has emerged independently in primate lentiviruses, gammaretroviruses, and Equine infectious anemia virus. The need of antagonizing SERINC5 has therefore shaped the genomes of diverse retroviruses.

Despite the absence of sequence similarity, S2, Nef, and glycoGag, share two crucial features: the presence of putative motifs capable of recruiting clathrin adaptors(15), and their ability to localize at the cell membrane. To counteract SERINC5, S2 requires an ExxxLL motif, which is predicted to interact with the clathrin adaptor AP2. ExxxLL is not conserved in all S2 alleles as it is absent in the EIAV^LIA^lineage (Fig. 4D). However, in this case ExxxLL is substituted with YxxL, making the canonical motif capable of engaging with the clathrin adaptor complex a fundamental feature of S2. As already shown for Nef(18), our results confirm that these infectivity factors may function as “adaptors of adaptors”, capable of connecting a targeted cargo (such as CD4(19), MHC-I(20) or SERINC5 and SERINC3) to clathrin adaptors. This model implies also the ability of Nef, glycoGag and S2 to interact with SERINC5 and SERINC3, which remains to be experimentally proven.

We have shown that S2, like Nef, can be myristoylated. As observed with Nef(4), myristoylation is essential for S2 counteraction of SERINC2. Recruitment of S2 at the plasma membrane could therefore be a prerequisite to facilitate the interaction with AP2 and to recruit SERINC5, following a model described in detail for Nef and CD4(19). Intriguingly, a canonical myristoylation signal, predicting Ser^6^ or Thr^6^ in addition to Gly^2^, is absent at the N-terminus of S2, making it an unusual substrate for the human N-myristoyl transferase, similar to other proteins, such as the cAMP-dependent kinase (21) and another retroviral protein like MLV Gag(22).

Surprisingly, irrespectively of S2, EIAV infectivity is only minimally affected by SERINC5. Such increased resistance to the host factor depends on EIAV Env, reminiscent of the glycoproteins derived from some HIV-1 isolates and from RNA viruses such as VSV and Ebola. EIAV Env has already been reported to promote virus entry via a pH-dependent pathway, therefore reinforcing the hypothesis that the endosomal uptake could provide a port of entry less sensitive to SERINC5 inhibition(4). It remains now to be established whether resistance to SERINC5 is a general feature shared by all EIAV isolates. Nevertheless, the question remains: why did EIAV, as HIV, apparently evolve redundant mechanisms to bypass inhibition by SSERINC5? We observe that this would not make a unique example of antagonist redundancy, since HIV-1 VpU and Nef, for example, have evolved complementary abilities to interfere with the cell surface level of the HIV-1 receptor CD4(23). Redundancy could therefore be, in some cases, necessary. Alternatively, it remains possible that, in addition to the effect on retrovirus infectivity, SERINC5 could exert other activities, similarly adverse to virus replication, which are not counteracted by Env. BST2, for example, was found to act both as a virus tethering factor, which prevents the release of virion particles in the extracellular space, and as a viral sensor, which signals through NFkB the presence of budding virus particles(24).

In conclusion, the identification of a third example of retroviruses, which independently evolved the SERINC5 and SERINC3 counteracting action, demonstrates that these host factors play a crucial inhibitory activity relevant to diverse retroviruses infecting different hosts. Altogether, this highlights a novel host-pathogen interface which should be investigated further to explore its utility for the development of innovative antiviral strategies.

## Materials and methods

### Plasmids

For all experiments performed with HIV-1, NL4-3-derived plasmids carrying a frameshift mutation to disrupt expression of *env* and/or *nef* were used as already described(25). Env-defective viruses were complemented with a PBJ5 plasmid expressing HIV-1^HXB2^env. Vectors FBASA was used to express Amphotropic (4070-A) MLV Env(26) and pMD.G to express the vesicular stomatitis virus G protein(27). EIAV codon optimized envelope pLGcoSUTM was constructed by inserting codon-optimized SU and TM sequences from pSPEIAV19 (GenBank accession number EIU01866) into the low-copy-number plasmid pLG228/30, as previously described(28). pCXCR4-FLAG has been described earlier(29). An *env*-defective NL4-3 expressing S2 in place of Nef was also generated by substituting the first 73 codons of *nef* with the coding sequence from S2 derived from SPEIAV19 (GeneBank accession n. U01866)(11). Codon-optimized version of S2 from SPEIAV19 and from pEV53D(30) (GeneBank accession n. M87581, derived from Wyoming strain EIAV) were obtained through a seven-step PCR, to substitute the required bases following human codon usage and cloned into PCDNA3.1. An HA tag was also added at the C-terminus by PCR. All S2 mutants with a HA tag fused at the C-terminus (L27,28AA, G2A, P26,29AA) were generated by site-directed mutagenesis. SERINC5-HA was expressed from pBJ6 (a derivative of PBJ5 which provides low level of expression in HEK 293T cells) and pSERINC5-GFP vectors were previously described(4). mRFP-Rab7 was a gift from Ari Helenius (Addgene plasmid # 14436)(31). EIAV transfer vector pEIAV-SIN6.1 CGFPW(32) and gag-pol expressing construct pEV53D were a gift from John Olsen (Addgene plasmid # 44171, # 44168 respectively). pEV53D in which S2 ORF was disrupted was obtained by enzymatic digestion of a unique BamHI site followed by Klenow fill-in and ligation, to insert a frameshift disrupting the S2 ORF from codon 18. Equus SERINC5 (accession XM_001503874) was custom-synthetized (GeneArt, Life Technologies) with codons optimized for human usage. AP180 C-terminus transdominant mutant was a gift from Harvey McMahon, LMB-MRC Cambridge, UK(12). Dynamin2 K44A, Nef G2A, hCypA, PBJ5 (28) Nef-HA were used as described earlier(25).

### Cell lines

See SI Materials and Methods

### Viruses and infectivity assay

HIV-1 limited to a single round of replication or EIAV vectors were produced by transfection of JTAg and CEM174X cells (electroporation) or HEK293T cells (the calcium phosphate) as described in SI materials and methods. Viruses were quantified using the SG-PERT reverse transcription assay(33) modified as described previously(34), diluted 3- or 5-fold in a series of 6 steps and used to infect TZM-zsGreen reporter cells Infectivity was assayed as a function of zsGreen-positive cells scored using the High Content Imaging System Operetta (Perkin Elmer) Infectivity was calculated by dividing the number of infected cells in a well for the amount of RT-activity associated to the virus inoculum, measured in mU(33). When indicated, results were expressed as percentage of an internal control sample. For a detail description of the procedure see SI Materials and Methods.

### Immunofluorescence and Western blotting

Standard procedures were used. For details refer to SI

### Myristic acid, azide pulse labelling and conjugation

HEK293T cells were pulsed with 50µM myristic acid-azide (Life Technologies) for 24 hours and the incorporated azide moiety was conjugated to the alkyne immobilized on a matrix using the Click-iT protein enrichment kit (Life Technologies), as reported earlier(35). HA-tagged proteins in the enriched fraction were detected by immunoblotting. For a detail description of this procedure see SI Materials and Methods.

## Acknowledgements.

We thank the Centre for AIDS Reagents, NIBSC, and NIH AIDS Research and Reference Reagent Program, Division of AIDS, for cell lines and plasmids. This work was supported by FP7 Marie Curie Career Integration Grant nr. 322130 and Caritro “Ricerca Biomedica” grant n. 2013.0248 to MP. HHS NIH R01CA128568 to SC.

